# PIEZO2 mediates ultrasonic hearing via cochlear outer hair cells in mice

**DOI:** 10.1101/2020.10.09.332643

**Authors:** Jie Li, Shuang Liu, Chenmeng Song, Qun Hu, Zhikai Zhao, Tuantuan Deng, Yi Wang, Tong Zhu, Linzhi Zou, Shufeng Wang, Jiaofeng Chen, Lian Liu, Hanqing Hou, Kexin Yuan, Hairong Zheng, Zhiyong Liu, Xiaowei Chen, Wenzhi Sun, Bailong Xiao, Wei Xiong

## Abstract

Ultrasonic hearing is exploited for hunting and navigation as in echolocation by microbats and bottleneck dolphins, and for social communication like ultrasonic vocalization by mice and rats. However, the molecular and cellular basis for ultrasonic hearing is not known yet. Here we show that knockout of the mechanosensitive ion channel PIEZO2 in cochlea disrupts the ultrasonic hearing but not the low-frequency hearing in mice, as shown by audiometry and acoustically-associative freezing behavior. Deletion of *Piezo2* in the outer hair cells specifically abolishes the associative learning of the mice upon hearing the ultrasonic frequency. *Ex vivo* cochlear Ca^2+^ imaging revealed that the ultrasonic transduction requires both PIEZO2 and the hair-cell mechanotransduction channel. Together, our study demonstrates that the outer hair cells are the effector cells with PIEZO2 as an essential molecule for ultrasonic hearing in mice.

**Significance Statement:** Some animals have evolved an incredible ability for vocalizing and hearing ultrasonic frequencies that is inaudible for humans (> 20 kHz). For many years, it has been considered that animals hear ultrasonic frequencies with their cochlear hair cells, using the identical set of mechanotransduction molecules in the hair bundles for hearing audible frequencies. Here, we show that the mice lacking the mechanosensitive ion channel PIEZO2 hardly hear ultrasonic frequencies, while can still be sensitive to audible frequencies. Thus, animals may use a partially different mechanism for sensing physiological ultrasound.

## Main Text

### Introduction

Some animals use ultrasonic hearing and vocalization to communicate and navigate in daily lives (1). For example, mice vocalize at frequencies > 25 kHz with intensities from 60 to 100 dB SPL during certain social behaviors, including mother-pup interaction, male-male encounter, and male-female courtship (2-4). Thus, the ultrasound-based auditory communication is critical for the survival and reproduction of mice. Studies on animal models, including mice, bats, cats, and guinea pigs, have provided neurophysiological insights into ultrasonic hearing (5-9). However, the precise molecule identity and cell-type definition concerning ultrasonic transduction are still elusive. Currently, ultrasonic hearing is often thought to share the general molecular and cellular mechanisms of auditory transduction that have been well recognized (10-13), which however is extremely putative for lack of experimental evidence.

It has been recently reported that the mechanosensitive ion channel PIEZO2 plays critical roles in the somatosensory system, including gentle touch, itch, tactile pain, proprioception, breath, and blood pressure (14, 15). Structural and functional analyses of PIEZO2 have shown that it can respond to different forms of mechanical stimuli, such as indentation and stretching (16, 17). Interestingly, PIEZO2 was found expressed at the apical surface (also known as cuticular plate) of the cochlear hair cells, mainly outer hair cells (OHCs) (18), to mediate a stretch-activated current (also known as reverse-polarity current) in neonatal mice (18, 19). However, this current gradually reduces with age and finally disappears around P7 (19, 20), which is opposite to the maturation of hair-cell mechanotransduction current (19, 21). Knockout of *Piezo2* in the inner ear only slightly affects hearing from 8 kHz to 20 kHz in adult mice as tested by the auditory brainstem response (ABR) recording (18). To date, the virtual physiological role of PIEZO2 in hearing is still elusive (20).

In this study, we explored the role of PIEZO2 in ultrasonic hearing from a variety of knockout (KO) and conditional knockout (cKO) mouse lines, using ultrasonically-combined approaches, including ABR recordings, behavior tests, and *ex vivo* cochlear imaging assays. We found that the expression of PIEZO2 in the OHCs is essential for ultrasonic hearing in mice.

## Results

### The mechanosensitive channel PIEZO2 is required for ultrasonic hearing

To evaluate ultrasonic hearing physiologically, we improved the basic ABR recording by connecting the electrode to a microscrew nailed at the skull bone positioned posterior to Bregma sutures (−7 mm AP, 0 mm ML) (Fig. 1A, Fig. S1 and Methods), named nail ABR (nABR) recording. The nABR configuration enhanced the detection stability and sensitivity to the stimuli at frequencies > 12 kHz in C57BL/6 (B6) mice (Fig. 1B and C). Although the ultrasonic responses were not as strong as those induced by low frequencies, the nABR waveforms induced by ultrasonic frequencies were distinguishable for determining the thresholds (Fig. 1B). The generally decreased nABR amplitude at 63 kHz and 80 kHz implies less efficient ultrasonic transduction at the cochlear level, because the ABR waveforms reflect signals from the auditory nerves that innervate the cochlear hair cells, and their ascending auditory pathways (22). The 54-kHz 90 dB SPL nABR signal suddenly showed a large amplitude (Fig. 1B), which consists with the fact that the mouse hearing has two peaked sensitivities at 15 and 55 kHz as previously reported by audiometry (23) and auditory nerve recordings (24). This phenomenon was not due to distortions delivered by the speaker at high intensities, since the measured ultrasonic pure-tone output was very condensed even at 90 dB SPL (Fig. S1D).

**Fig 1.**
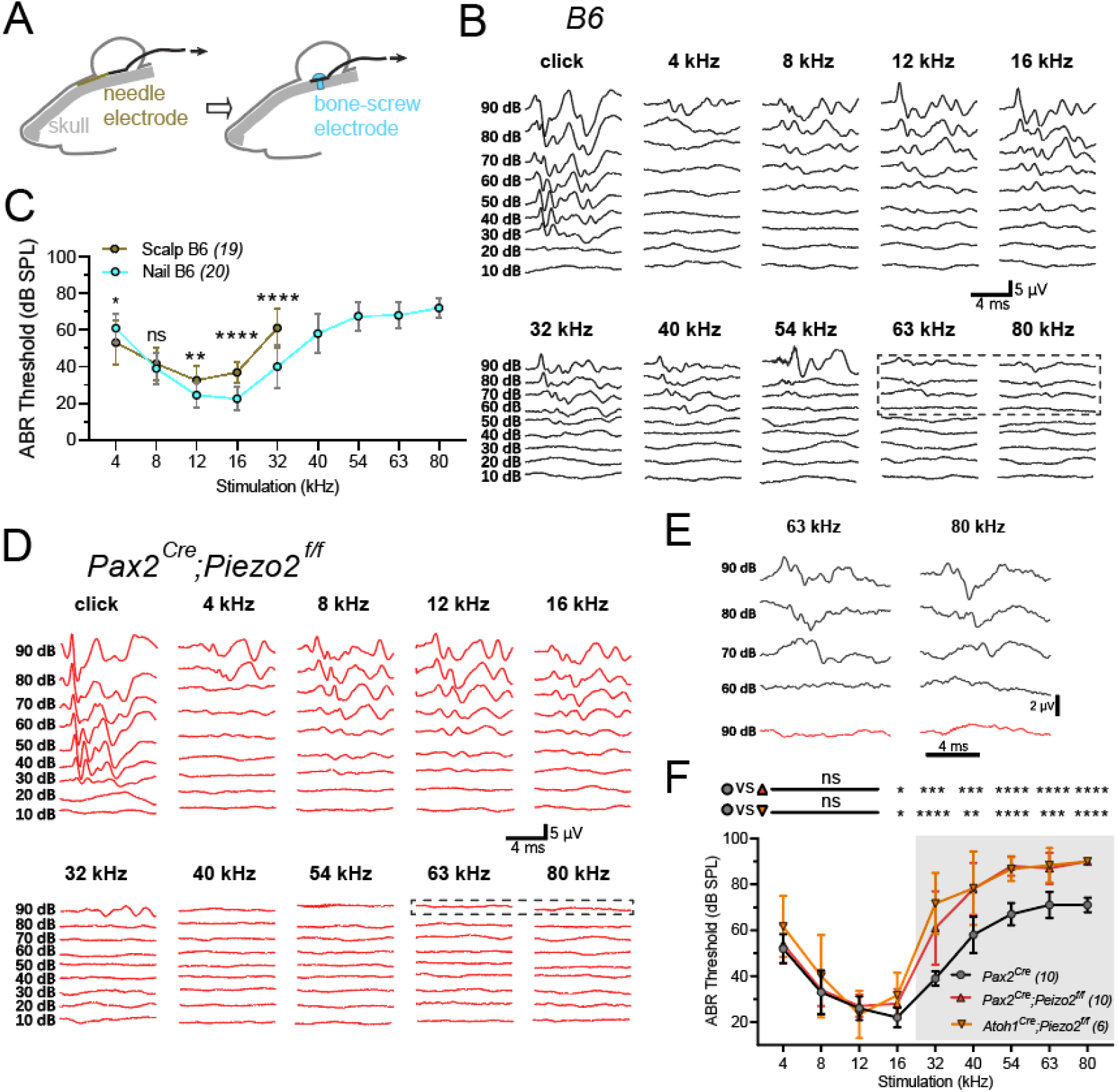
The mechanosensitive channel PIEZO2 is required for ultrasonic hearing. (*A*) Schematic of recording configuration of nABR. For nABR, the recording wire was connected with a stainless-steel bone screw implanted on the mouse skull (cyan) instead of placing a needle electrode under scalp (dark yellow) (see fig. S1). (*B*) Representative example of nABR signals in a C57BL/6J (B6) mouse. (*C*) Comparing to regular ABR in B6 mice with electrode under scalp (Scalp B6, dark yellow), the nABR achieved an improved sensitivity to frequencies > 12 kHz in B6 mice (Nail B6, cyan). Scalp B6 vs Nail B6, unpaired *t*-test, 4 kHz, *p = 0.02; 8 kHz, p = 0.36 (no significance, ns); 12 kHz, **p < 0.002; 16 kHz, ****p < 0.0001; 32 kHz, ****p < 0.0001; error bars, SD. (*D*) Representative example of nABR signals in *Piezo2*-cKO mouse. (*E*) Enlarged traces with 63 kHz and 80 kHz sound stimuli framed in (*B*) and (*D*). (*F*) Pure-tone nABR thresholds in control *Pax2*^*Cre*^ mice and *Piezo2*-cKO mice. The control mice and *Piezo2*-cKO mice showed distinct ABR thresholds on ultrasonic frequencies (gray-shaded area). *Pax2*^*Cre*^ mice vs *Pax2*^*Cre*^;*Piezo2*^*f/f*^ cKO mice, Unpaired *t*-test, 4 kHz, p = 0.556; 8 kHz, p = 0.791; 12 kHz, p = 0.66; 16 kHz, *p = 0.022; 32 kHz, ***p = 0.0005; 40 kHz, ***p = 0.0002; 54 kHz, ****p<0.0001; 63 kHz, ****p<0.0001; 80 kHz, ****p<0.0001; error bars, SD. *Pax2*^*Cre*^ mice vs *Atoh1*^*Cre*^;*Piezo2*^*f/f*^ cKO mice, unpaired *t*-test, 4 kHz, p = 0.0669; 8 kHz, p = 0.319; 12 kHz, p = 0.4985; 16 kHz, *p = 0.0153; 32 kHz, ****p<0.0001; 40 kHz, **p = 0.0041; 54 kHz, ****p<0.0001; 63 kHz, ***p<0.0002; 80 kHz, ****p<0.0001; error bars, SD. All the mice were used at the age about 1 month. For (*C*) and (*F*), N numbers are shown in panels.

As the *PIEZO2* pan-knockout mice are embryonic-lethal (25), we crossed certain Cre mice with *Piezo2*^*f/f*^ mice (26) to generate *Piezo2-*cKO mice, which contained mixed genetic background. To avoid possible influence of the genetic background on evaluation of ultrasonic hearing, we compared the nABR sensitivity between the B6 mice and the CBA/J (CBA) mice. It has been shown that the CBA mice have better ultrasonic hearing sensitivity compared to the B6 mice according to the auditory nerve recordings (8-16 weeks animals) (24) and ABR recordings (16-18 weeks animals) (27), and the B6 mice exhibit progressive hearing loss late in life (> 7 months) (28, 29). Thus we recruited the B6 mice and the CBA mice at the age around 1 month when they have had matured auditory function while before age-related hearing loss may start to occur. The nABR recordings show that the ultrasonic hearing of the B6 mice is as sensitive as that of the CBA mice at the age of 1 month (Fig. S1E). In addition, the male mice and the female mice showed similar nABR thresholds (Fig. S1F). Next we examined the hearing and auditory transduction on the hybrid *Piezo2-*cKO mice with their littermates used as control at the age around 1 month, unless otherwise stated.

To check whether PIEZO2 participates ultrasonic hearing, we compared the nABR of the inner-ear targeted *Pax2*^*Cre*^*;Piezo2*^*f/f*^ mice (18, 30) and the cochlea targeted *Piezo2*-cKO mice by crossing the *Piezo2*^*f/f*^ mice with the *Atoh1*^*Cre*^ mice (31). The control mouse showed notable nABR signals at 32-80 kHz (Fig. 1B), while the *Pax2*^*Cre*^*;Piezo2*^*f/f*^ cKO mouse showed decreased response at frequencies > 32 kHz (Fig. 1D). This difference can be clearly seen when comparing their ABR responses together (Fig. 1E). The summarized nABR recordings reveal that the two types of *Piezo2*-cKO mice both had significantly reduced sensitivity of ultrasonic hearing (16-80 kHz) specifically but not to low-frequency hearing (4-12 kHz) (Fig. 1F). These data thus demonstrate that PIEZO2 is required for ultrasonic hearing.

We next examined whether lack of ultrasonic hearing is due to loss of the “high-frequency” hair cells at the very basal coil of cochlea in the *Piezo2*-cKO mice. No obvious loss of hair cells was found in the cochleae of the *Pax2*^*Cre*^*;Piezo2*^*f/f*^ cKO mice, which preserved normal morphology of the hair cells (Fig. S2). We further examined the organization of the inner ear of the *Pax2*^*Cre*^*;Piezo2*^*f/f*^ cKO mouse by a tissue clearing approach (32) (Methods). The whole structure of the inner ear was intact and the hair cells kept normal allocation and abundance (Movie S1 and S2).

### Ultrasonically-associative freezing behavior is disrupted in *Piezo2*-knockout mice

Next, we wondered whether PIEZO2-mediated ultrasonic hearing sensitivity is required for learned behavior in animals. The hearing response of the *Piezo2*-cKO mice was examined by a fear conditioning test (Fig. 2A), which associates an acoustic cue to the freezing behavior after paired training of the acoustic cue with electrical shocks on mice. An 90-dB SPL ultrasonic 63-kHz stimulatory cue was used because the 63-kHz nABR showed a stable threshold difference between the control and the *Piezo2*-cKO mice (Fig. 1F), and 63 kHz is in the range of ultrasonic frequencies of mouse social communications (1). To exclude the possibility that the disrupted ultrasonically-associative fear conditioning test was due to learning defect in *Piezo2*-cKO mice, freezing behavior with 16 kHz 90 dB SPL cue was performed as a control, with the same experimental set up. Harmonics appears in the 16-kHz stimulation but with intensities lower than the ABR threshold (Fig. S3A). The sound intensity measured near the arena floor was from 75 dB SPL to 95 dB SPL (Fig. S3B), which is larger than the ultrasonic hearing threshold of mice. We also examined the locomotion activity of tested mice with different genotypes in an open field. The control and Piezo2-cKO mice showed no difference in locomotion distance in 5 minutes statistically (Fig. S3C).

**Fig 2.**
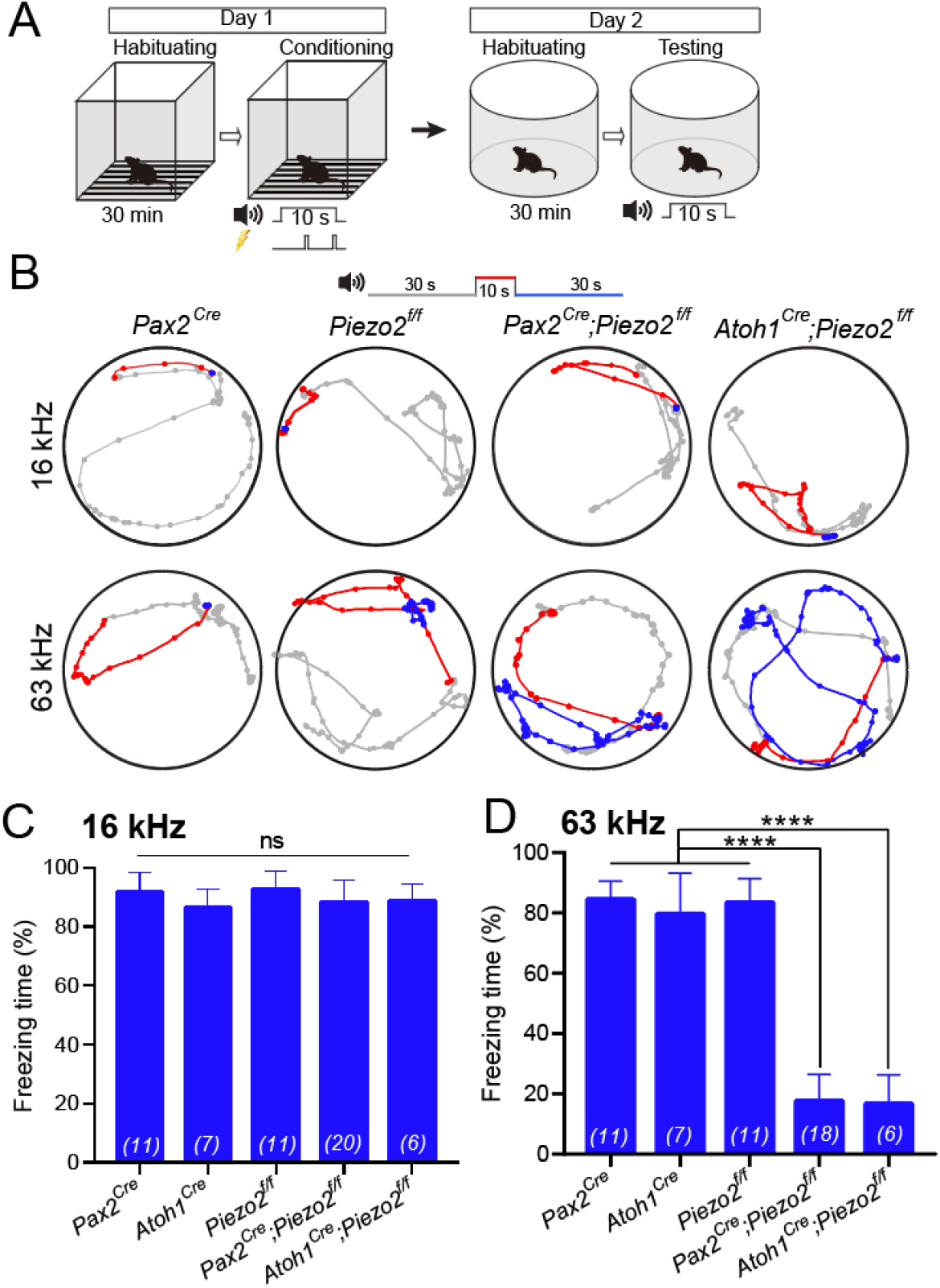
Ultrasonically-associative freezing behavior is disrupted in *Piezo2*-cKO mice. (*A*) Paradigm of sound-cue associated freezing behavior. Pure-tone sound at 16 kHz or 63 kHz played by a TDT ES1 (Free Field) electrostatic speaker was used as the conditional stimulation, and footshock was used as the unconditional stimulation. (*B*) Representative examples of locomotion of control mice and *Piezo2*-cKO mice before (gray, 30 s), during (red, 10 s), and after (blue, 30 s) the pure-tone sound cue at 90 dB SPL. The mice had been trained to pair either 16-kHz cue or 63-kHz cue with the footshock-induced freezing. Dots indicate the location of mouse every 0.5 s. *Pax2*^*Cre*^ mice, *Atoh1*^*Cre*^ mice and *Piezo2*^*f/f*^ mice were used as controls. (*C* and *D*) Freezing time in percentage with 16-kHz cue (*C*) or 63-kHz cue (*D*). In (*D*), two *Pax2*^*Cre*^;*Piezo2*^*f/f*^ mice were omitted because they completely had not locomotion during test. One-way ANOVA, p = 0.225 in (*C*), ****p < 0.0001 in (*D*); error bars, SD. N numbers are shown in panels. All the mice were about 1 month old.

Comparison of freezing behavior to the 90-dB 16-kHz cue for the control *Pax2*^*Cre*^ mice and *Piezo2*^*f/f*^ mice with the *Pax2*^*Cre*^*;Piezo2*^*f/f*^ and *Atoh1*^*Cre*^*;Piezo2*^*f/f*^ cKO mice showed that all genotypes retained their low frequency-associative freezing behavior (Fig. 2B), as shown in freezing time percentage (Fig. 2C), indicating that these mice all possess low-frequency hearing and acoustically-associative learning. In contrast, as for 63-kHz cue, the *Pax2*^*Cre*^*;Piezo2*^*f/f*^ and *Atoh1*^*Cre*^*;Piezo2*^*f/f*^ cKO mice showed disrupted freezing behavior (Fig. 2B and D), while this ultrasonically learned behavior was preserved in the control mice (Fig. 2D). These data show that PIEZO2 is required for mice to behaviorally respond to ultrasound.

### Expression of PIEZO2 in cochlear hair cells

Next we examined the expression of PIEZO2 in the cochleae. Firstly, the *Piezo2-GFP-IRES-Cre* mice that carry a Cre cassette with the endogenous *Piezo2* (26) were crossed with the *H2B-mCherry* mice that have Cre-inducible mCherry expression (33). The mCherry expression was observed in most OHCs and some inner hair cells (IHCs) in 1-month mice (Fig. S4A-D) as previously reported (18), indicating *Piezo2* promoter is transcriptional active in cochlear hair cells. Due to lack of suitable PIEZO2 antibody, we used a GFP antibody to locate PIEZO2 expression in the *Piezo2-GFP-IRES-Cre* mice that has a GFP gene fused with *Piezo2*. PIEZO2 immunostaining signal was mainly detected at the apical surface of cochlear OHCs by the GFP antibody at postnatal day 5 (P5) (Fig. 3A) as reported (18), which however was hardly detectable at more mature age, e.g. 3-4 weeks. Thus we applied a RNAscope protocol (34) to check the expression of *Piezo2* transcript in control and *Pax2*^*Cre*^*;Piezo2*^*f/f*^ mice at P21. Within the cross-section of organ of Corti, *Piezo2* transcript was observed in the control *Piezo2*^*f/f*^ hair cells though at a relatively low level, while the transcript was significantly reduced in *Pax2*^*Cre*^*;Piezo2*^*f/f*^ hair cells (Fig. 3B and C). Control staining in either the inner ears or the dorsal root ganglia was performed to validate our optimized RNAscope procedure is effective (Fig. S4E and F). These data indicate that PIEZO2 is expressed in the cochlear hair cells of adult mice.

**Fig 3.**
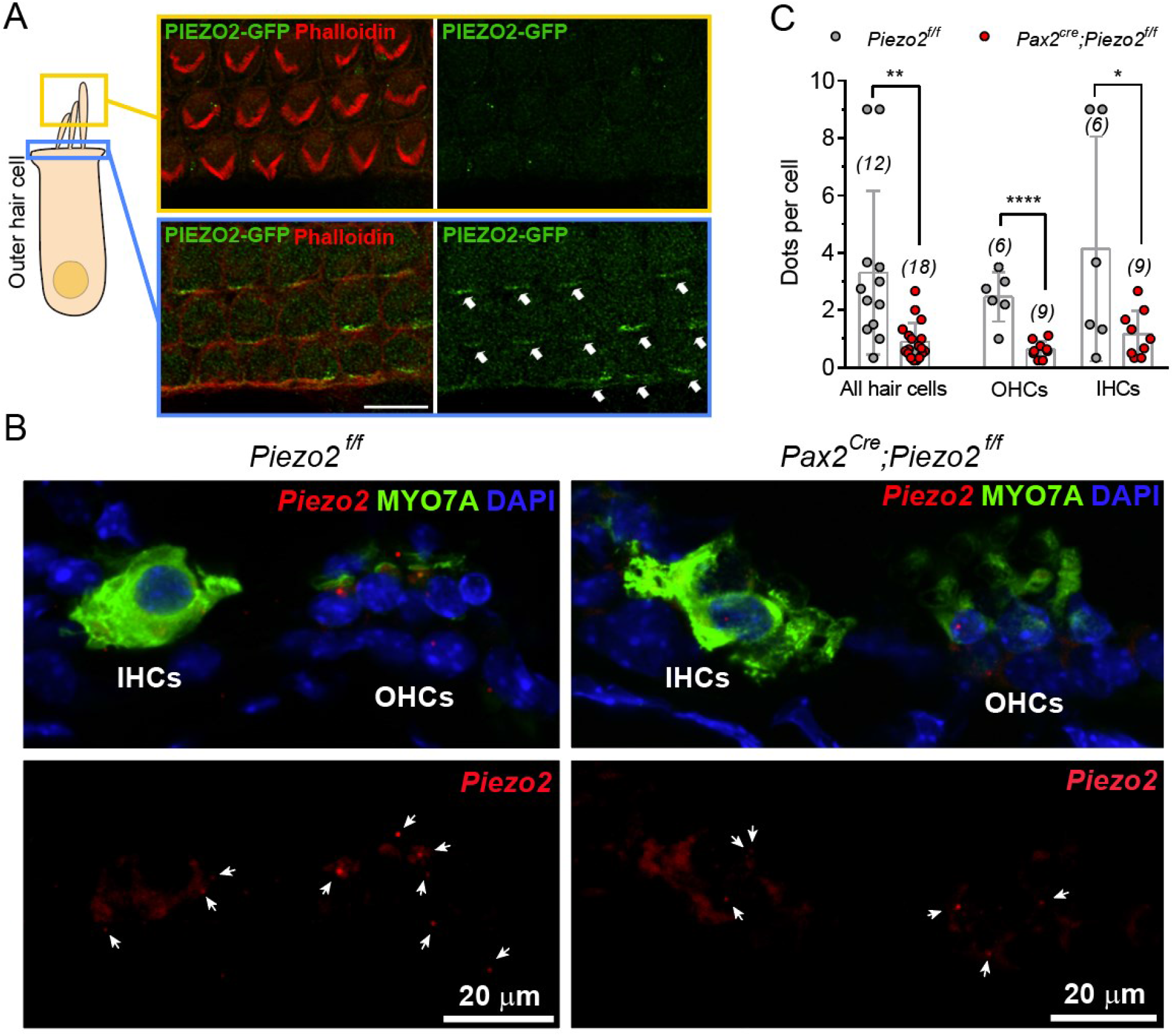
Expression of Piezo2 in cochlear hair cells. (*A*) Schematic showing hair bundle layer (yellow) and cuticular plate (blue) of an outer hair cell. Piezo2 signal (green) was detectable at cuticular plate (lower panel in blue frame, white arrows) but not in hair bundles (upper panel in yellow frame), as stained by GFP antibody (green) in P5 *Piezo2-GFP* mice. Hair bundle was stained by Phalloidin (red). Bar, 20 μm. (*B*) Cross-sections of organ of Corti of a *Piezo2*^*f/f*^ mouse and a *Pax2*^*Cre*^;*Piezo2*^*f/f*^ mouse at P21 showing fluorescent signals of RNAscope probe targeting *Piezo2* (red dots, indicated by arrows). Some dots are not clear due to out of focus. Hair cells and nuclei were stained by MYO7A antibody (green) and DAPI (blue), respectively. Bars, 20 μm. (*C*) Quantification of *Piezo2* dots in cochlear hair cells from data similar to (*B*). Cochlear sections were collected from P21 mice for each genotype. Note there are multiple hair cells superimposed. Numbers of transcript dots were counted per section and numbers of hair cells were counted based on MYO7A and DAPI signals. Unpaired *t*-test, All hair cells, **p < 0.002; OHCs, ****p < 0.0001; IHCs, *p = 0.043; error bars, SD; N numbers of sections are shown in panels.

### Mice lack ultrasonic hearing when *Piezo2* was deleted in outer hair cells

We further wondered which type of hair cell supports PIEZO2’s role for ultrasonic hearing by examining the ultrasound-associative freezing behavior in mice when deleting *Piezo2* in OHCs or IHCs. The *Prestin*^*CreER*^ mice (35) was introduced to generate OHC-specific *Piezo2*-cKO mice as *Prestin* only expresses in OHCs, and the *vGlut3*^*CreER*^*;Piezo2*^*f/f*^ mice were used to check PIEZO2’s role in IHCs (36). To validate the hair-cell expression specificity, the *Prestin*^*CreER*^*;H2B-mCherry* mice and the *vGlut3*^*CreER*^*;H2B-mCherry* mice were injected with tamoxifen at P8-10, a time with peaked expression of *Prestin* and *vGlut3*, and later examined for the mCherry expression at 1-2 month age (Fig. S5A). Only OHCs showed mCherry expression with a high efficiency in the *Prestin*^*CreER*^*;H2B-mCherry* mice (Fig. S5B), while mCherry was widely expressed in IHCs of the *vGlut3*^*CreER*^*;H2B-mCherry* mice (Fig. S5C). With the same injection procedure (Fig. 4A), the induced *Prestin*^*CreER*^*;Piezo2*^*f/f*^ cKO mice at 1 month showed freezing behavior with the low-frequency stimulation but not the ultrasonic cue (Fig. 4B and C), indicating a necessity of OHCs for ultrasonic hearing. On the contrary, the *vGlut3*^*CreER*^*;Piezo2*^*f/f*^ mice did not show any deficit of ultrasound or low-frequency sound associative freezing (Fig. 4D and E), which excludes a possible role of IHCs in PIEZO2-mediated function in ultrasound detection. These data confirm that the expression of PIEZO2 in cochlear OHCs is required for ultrasonic hearing.

**Fig 4.**
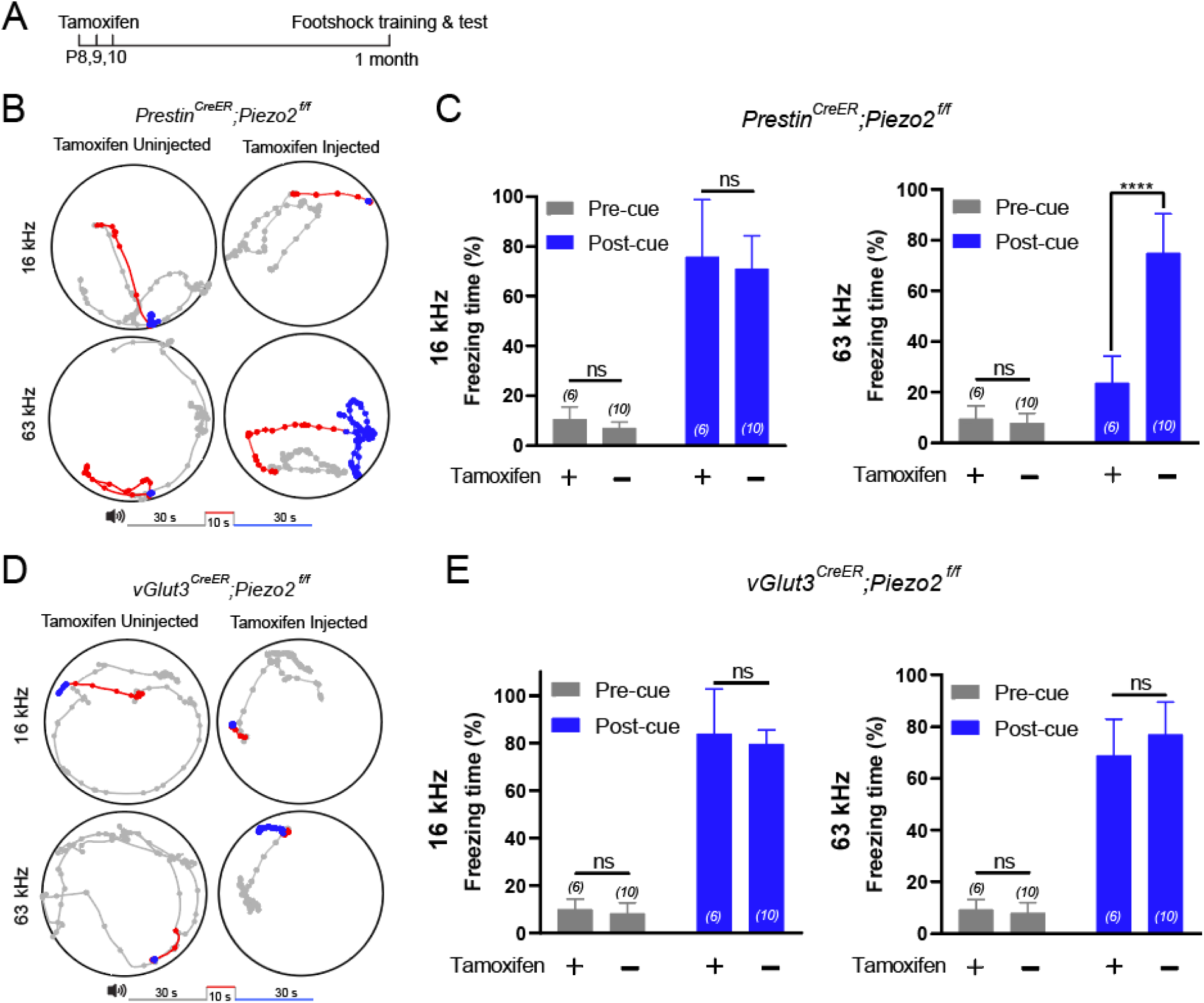
Mice lack ultrasonic hearing when *Piezo2* was deleted in outer hair cells. (*A*) Schedule of tamoxifen injection and behavior test. (*B*) Representative example of locomotion of *Prestin*^*CreER*^;*Piezo2*^*f/f*^ mice with or without tamoxifen injection. Different colors represent the locomotion before (gray), during (red), and after (blue) the 16-kHz or 63-kHz sound cue. Dots indicate the location of mouse every 0.5 s. (*C*) Freezing time (in percentage) of the *Prestin*^*CreER*^;*Piezo2*^*f/f*^ mice trained and tested with 16-kHz cue (left panel) or 63-kHz (right panel) cue. Unpaired *t*-test, 16 kHz, pre-cue, p = 0.202; post-cue, p = 0.669; 63 kHz, pre-cue, p = 0.424; post-cue, ****p < 0.0001; error bars, SD. (*D*) Representative example of locomotion of *vGlut3*^*CreER*^;*Piezo2*^*f/f*^ mice with or without tamoxifen injection. Other conditions are the same with (*B*). (*E*) Freezing time of the *vGlut3*^*CreER*^;*Piezo2*^*f/f*^ mice trained and tested with 16-kHz cue (left panel) or 63-kHz (right panel) cue. Unpaired *t*-test, 16 kHz, pre-cue, p = 0.428, post-cue, p = 0.492; 63 kHz, pre-cue, p = 0.174, post-cue, p = 0.203; error bars, SD. For (*C*) and (*E*), N numbers are shown in panels.

### Hair-bundle mechanotransduction is required for ultrasonic transduction

We next questioned whether the hair-bundle mechanotransduction participates in the ultrasonic transduction. It has been reported that TMC1 is the putative auditory transduction channel in hair cells and CDH23 is one of the two tip-link components, so we used the *TMC1*-KO mice (37) and the CDH23-null *v2j* mice (38) to investigate their ultrasonic transduction. However, the two mutant mice had complete loss of hearing from low frequencies to ultrasonic frequencies as shown by nABR recordings at 1 month age (Fig. S6A), likely resulted from abnormal hair bundles (37, 38) and loss of the hair cells (Fig. S6B and C). These results prompt that audiometry or behavior is not proper to probe the contribution of hair-bundle mechanotransduction and an approach with cellular resolution is needed.

To directly monitor the ultrasonic transduction in the cochlear hair cells, we customized an *ex vivo* ultrasonic stimulation stage delivering ultrasonic vibration of 80 kHz - a frequency within the range of the physiological hearing of mice (Fig. S7A-C and Methods). This stimulation mimics the mechanical vibration in the cochlea driven by incoming ultrasound. It is difficult to obtain the organ of Corti from mice after hearing onset because the cochlea has been embedded into the bony capsule of the inner ear. Instead, we introduced the hemicochlear preparation (39, 40) that preserves most of the elements of the cochlea and is also accessible for microscopic observation (Fig. 5A). Because the patch-clamp recording of the hair cells was always destroyed by direct ultrasonic stimulation, we thus used Ca^2+^ imaging for monitoring the ultrasonically-evoked activity (Fig. 5B). The hemicochlear preparation was loaded with Fluo-8 AM, a sensitive Ca^2+^ dye, and OHCs were the major cells with Ca^2+^ dye uptake (Fig. 5C). The ultrasonic stimulation elicited Ca^2+^ waves in the OHCs of WT hemicochleae despite of the position (apical or middle) in the cochlear coil, which could be blocked when perfusing the 0.1 mM Ca^2+^ solution (Fig. S7D and E), prompting that the OHCs are ultrasonically-responsive. After ultrasonic stimulation, the Fluo-8 loaded OHCs could show evoked Ca^2+^ wave when ATP (100 μM) was applied (Fig. S7D and E), indicating that the OHCs were healthy after ultrasonic stimulation and the Ca^2+^ response was not saturated.

**Fig 5.**
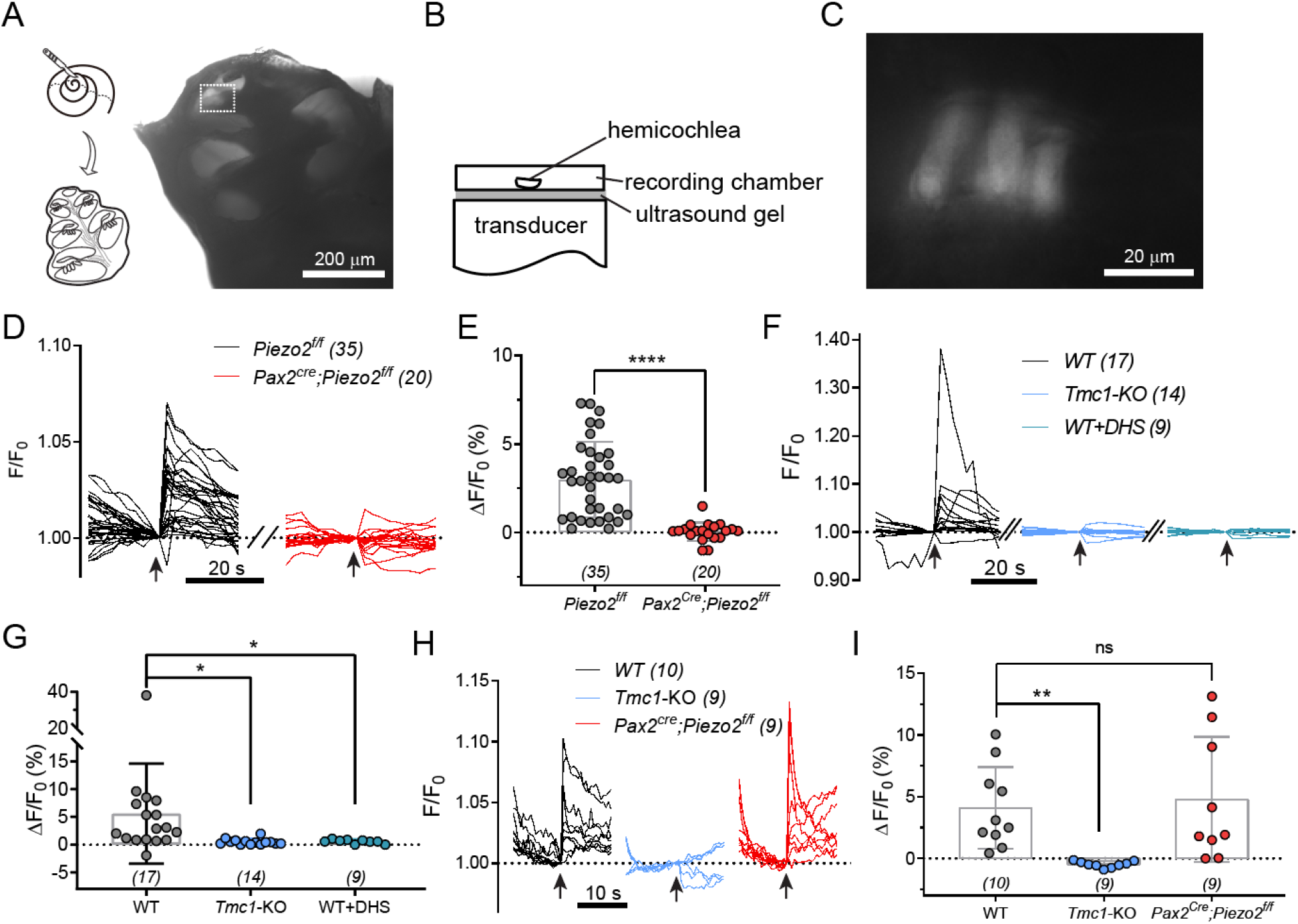
Hair-bundle mechanotransduction is required for ultrasonic hearing. (*A*) Left: schematic showing preparation of hemicochlea. Right: a photo of hemicochlea with transmission-light illumination. Bar, 200 μm. (*B*) Setup for ultrasonic transducer stimulated hemicochlea. An 80-kHz transducer was fixed underneath the recording dish with ultrasound gel in between. (*C*) A fluorescent image showing Fluo-8 AM loaded OHCs, magnified from the apical part (white-dashed frame) of the hemicochlea in (*A*). Bar, 20 μm. (*D*) Ultrasonic stimulation evoked Ca^2+^ responses of OHCs in hemicochlea preparations from control *Piezo2*^*f/f*^ mice (black) and *Pax2*^*Cre*^;*Piezo2*^*f/f*^ cKO mice (red). Arrows indicate the onset of ultrasonic stimulation. The images were collected at 2 s interval. See Movie S3 and S4. (*E*) Quantification of the peak Ca^2+^ responses of OHCs calculated from recordings in (*D*). Unpaired *t*-test, ****p < 0.0001; error bars, SD. (*F*) Ultrasonic stimulation evoked Ca^2+^ responses of OHCs from WT hemicochleae, *Tmc1*-KO hemicochleae, and WT hemicochleae treated with 100 μM DHS. (*G*) Quantification of the peak Ca^2+^ responses of OHCs calculated from recordings in (*F*). WT hemicochleae vs *Tmc1*-KO hemicochleae, unpaired *t*-test with Welch’s correction, *p = 0.033; error bars, SD. WT hemicochleae vs WT hemicochleae treated with 100 μM DHS, unpaired *t*-test with Welch’s correction, *p = 0.036; error bars, SD. (*H*) Low-frequency fluidjet evoked Ca^2+^ responses of OHCs in hemicochlea preparations from WT, *Tmc1*-KO and *Pax2*^*Cre*^;*Piezo2*^*f/f*^ cKO mice. (*I*) Quantification of the peak of OHCs in hemicochlea preparations from WT, *Tmc1*-KO and *Pax2*^*Cre*^;*Piezo2*^*f/f*^ cKO mice from similar recordings in (*H*). WT hemicochleae vs *Tmc1*-KO hemicochleae, unpaired *t*-test with Welch’s correction, **p = 0.0017; error bars, SD. WT hemicochleae vs *Pax2*^*Cre*^;*Piezo2*^*f/f*^ cKO hemicochleae, unpaired *t*-test, p = 0.733; error bars, SD. For (*E*), (*G*), and (*I*), N numbers are shown in panels. In this figure, all the mice were used at age about 1 month.

Next we checked the roles of PIEZO2 and TMC1 in ultrasonic transduction by hemicochlear Ca^2+^ imaging. The ultrasonic stimulation could elicit Ca^2+^ response in OHCs of the control *Piezo2*^*f/f*^ mice, but hardly in OHCs when genetically removing *Piezo2* (Fig. 5D and E, and Movie S3 and S4). Although the OHCs were widely lost in *Tmc1*-KO mice from 3 weeks (Fig. S8B) (37), we could still chose the apical OHCs with clear soma shape illuminated by Fluo-8 for Ca^2+^ imaging. The ultrasonic Ca^2+^ response was hardly seen in the *Tmc1*-KO OHCs or in the WT OHCs blocked by the mechanotransduction channel blocker dihydrostreptomycin (DHS, 100 μM) (Fig. 5F and G), implying a requirement of the hair-bundle mechanotransduction in the ultrasonic transduction. To investigate the response evoked by the low-frequency stimulation, we used fluid jet, a stimulation directly deflects the hair bundle, to induce the mechanotransduction channel mediated Ca^2+^ response in *Tmc1*-KO or *Pax2*^*Cre*^*;Piezo2*^*f/f*^ OHCs. Similar to ultrasonic frequencies, there is no fluid-jet evoked Ca^2+^ response in *Tmc1*-KO OHCs. In contrast, the fluid-jet induced Ca^2+^ response was maintained in *Pax2*^*Cre*^*;Piezo2*^*f/f*^ OHCs and WT OHCs (Fig. 5H and I). These *ex vivo* results showed that PIEZO2 is required for the OHC Ca^2+^ response stimulated by ultrasonic frequencies, but not for those stimulated by low frequencies. We further tested whether PIEZO2 and/or TMC1 could establish the ultrasonic transduction in exogenous expression systems. Although the HEK293T cells expressing PIEZO2 were mechanosensitive (Fig. S8A and B), we did not observe any response in the HEK293T cells expressing PIEZO2 or PIEZO2 and TMC1 when using the same 80-kHz ultrasonic stimulation applied for the hemicochlear imaging (Fig. S8C and D). These results show that PIEZO2 may coordinate with the hair-bundle mechanotransduction machinery to fulfil the ultrasonic transduction.

## Discussion

Here we show multiple lines of evidence that PIEZO2 is essential for mice to hear ultrasonic frequencies within the range necessary for social communication in mice. Furthermore, the ultrasonic transduction is mainly conducted by OHCs, which coordinates the hair-bundle mechanotransduction. Small mammals, such as mice, emit strong ultrasonic vocalization for their social communication while overriding the mask effect from ambient noise or not being heard by their predators. As the molecular and cellular mechanisms of ultrasonic hearing and transduction have not yet been clear, our work points out a possible mechanism which may be overlooked before.

Previous evidence has shown that at neonatal age, PIEZO2 is expressed in cochlear hair cells (18). Our findings further prompt that the PIEZO2 expression in OHCs continues to at least week 3-4 as examined (Fig. 3B and C). The *Piezo2* expression shown by RNAscope assay is few, which may not reflect the quantity of PIEZO2 in OHCs. It has been shown that with a low level of *Piezo1* expression, PIEZO1-mediated mechanically activated current could be detected in cell lines (41). PIEZO2’s mechosensitivity properties likely support its role for OHCs in ultrasonic transduction. PIEZO2 is a mechanosensitive channel that homotrimerizes to form a gigantic (∼0.9 Mega-Dalton) three-bladed propeller-like structure comprising 114 transmembrane (TM) domains (38 TM per protomer), making it a unique membrane protein with the largest number of TMs (16). Strikingly, the three unusual non-planar TM Blades are curved into a nano-bowl shape of 28 nm-diameter and 10 nm-depth, which might deform the residing membrane to produce a mid-plane nano-bowl surface area of 700 nm^2^ and a projected in-plane area of 450 nm^2^. On the basis of the unique nano-bowl shape of the Piezo channel-membrane system, flattening the non-planar TM-Blades might produce a maximal change of the projection area of ∼250 nm^2^, which might provide the energy to gate the channel (16). The curved configuration of the Piezo channels (PIEZO1 and PIEZO2) might further deform the membrane shape outside of the perimeter of the channel into a large, curved ‘membrane footprint’ (42), which might further amplify the mechanosensitivity of the Piezo channels. Such ‘membrane-dome’ (43) and ‘membrane footprint’ (42) mechanisms have been proposed to account for the exquisite mechanosensitivity of Piezo channels in response to various mechanical stimulation including poking and stretch, which may underlie the Piezo1’s response to the non-physiological ultrasonic stimulation (0.5 MHz) (44). However, removal of either PIEZO2 or TMC1 abolished ultrasonic transduction (80 kHz) in cochlear hair cells (Fig. 6, D to G), suggesting there is a more complicated mechanism for PIEZO2 to physiologically transduce or detect ultrasonic frequencies. PIEZO2 may locate at the apical surface of the hair cell, where the stereocilia root and is a place least influenced by the membrane low-pass filtering, and coordinate with the hair-bundle mechanotransduction machinery to transduce the ultrasonic vibration. Thus, it is possible that PIEZO2 itself may only detect the ultrasonic waves instead of transducing it. This also explains why the HEK293T cells expressing PIEZO2 and/or TMC1 failed to show ultrasonic response (Fig. S8). By establishing the hemicochlear imaging, we found that in this period PIEZO2 and the hair-bundle mechanotransduction are both required for ultrasonic transduction. We think that these data intriguingly reflect a functional consolidation of the two molecularly distinct mechanotransduction channels during hearing maturation. As Wu et al. proposed (18), the mechanosensitivity and Ca^2+^ modulation mediated by PIEZO2 may support the normal development of hair cells. In the first neonatal week, the hair-bundle structure and function are still in development, which may prevent the OHCs to correctly distinguish the two channels from their physiological function. Thus, during this time, the two mechanotransduction machinery are antagonized, e.g. disruption of stereociliary mechanotransduction would unmask PIEZO2-mediated mechanotransduction (18, 20). While after onset of hearing, the two mechanotransduction channels establish their physiological functions respectively – TMC1-based machinery for sonic transduction and PIEZO2-mediated mechanism for ultrasonic detection.

Interestingly, the cochlear OHCs, not the IHCs, are the effectors to support PIEZO2 to sense ultrasound (Fig. 4), by which the animal gains extended spectral sensitivity from 16 kHz toward ultrasonic frequencies (Fig. 1F). Another OHC-specific protein is Prestin that enhances the cochlear mechanics and hearing sensitivity by providing OHCs somatic electromotility. With PIEZO2, the OHCs may use a mechanism like ciliary motility to transfer the ultrasonic vibration to the IHCs through the relative motion between the tectorial membrane and the reticular lamina, which however needs further empirical evidence. The IHC may simply output the encoded ultrasound information from the organ of Corti, implied by the evidence that the induced *vGlut3*^*CreER*^*;Piezo2*^*f/f*^ mice have normal ultrasonic freezing (Fig. 4D and E).

A similar examination of *Piezo2*-null mice with ABR measurement was previously reported by Wu et al. (18) but in lower frequency range. In their paper, mild differences (< 10 dB) in ABR thresholds were observed in 12 – 28 kHz between *Piezo2*-cKO and *Pax2*^*Cre*^ mice, which are similar with our results (Fig. 1F). However, the observation that *Piezo2*-cKO and *Pax2*^*Cre*^ mice have similar ABR thresholds in 32 kHz seems odd with our findings. Also, in our study, the control *Pax2*^*Cre*^ mice have similar nABR threshold with the B6 mice (Fig. 1C and D), which is slightly different from their observation that the 2-month *Pax2*^*Cre*^ mice showed higher ABR thresholds (18). Several factors may contribute to these differences. First, the two studies used different ABR configurations. Using a screw electrode implanted in the skull while directly attached to the brain, the nABR configuration adapted in our study achieved a higher stability and sensitivity upon ABR signal, especially in ultrasonic hearing range. Second, we used 1-month old mice while they used 2-month old mice for ABR test. The hearing sensitivity at different ages might be slightly different. Third, different genetic background for generating hybrid mice may result in shifts in their physiological functionality. Our data show that ultrasonic hearing is established from P21 at least. Thus most of our recordings were collected from 1-month old mice with their littermate as control, which avoids potential progressive degeneration that results in misinterpretation.

In summary, we discovered that PIEZO2 in OHCs plays an indispensable role in the ultrasonic high-frequency hearing, suggesting an alternative auditory transduction mechanism for frequencies in mammals. Given that both ultrasonic hearing and low-frequency hearing are conducted via cochlear OHCs but may be based on different mechanisms, it will be interesting to investigate the responding pattern to ultrasonic frequencies in cochlea, i.e. whether it follows the place-code principle. It would be intriguing to study PIEZO2’s role in ultrasonic hearing from other species, e.g. bats and whales. Moreover, our study lays the foundation to further address whether ultrasonic hearing and low-frequency hearing use distinct neural circuits and processing principles in the brain regions along the ascending auditory pathways.

## Materials and Methods

### Mouse strains and animal care

In this study, *Cdh23*^*v-2j*^ mice (Stock No. 002552, named *Cdh23*-null in this study), B6.129-*Tmc1*^*tm1*.*1Ajg*^/J mice (Stock No. 019146, named *TMC1*-knockout in this study), and *Rosa26 LSL H2B mCherry* mice (Stock No. 023139, named *H2B-mCherry* in this study) were from the Jackson Laboratory (Bar Harbor, ME); *Pax2-Cre* mouse line (named *Pax2*^*Cre*^ in this study) was generated by Dr. Andrew Groves (30) and *Atoh1-Cre* mice (named *Atoh1*^*Cre*^ in this study) were kindly from Dr. Lin Gan (31); *Piezo2*^*loxP/loxP*^ (named *Piezo2*^*f/f*^ in this study) and *Piezo2-GFP-IRES-Cre* mice were gifts from Dr. Ardem Patapoutian at the Scripps Research Institute (26); Prestin-CreER^T2^ (*Prestin*^*CreER*^ in this study) mouse line was a gift from Dr. Jian Zuo (35); *vGlut3-P2A-iCreER* knockin mouse strain was generated as described (36) and here named as *vGlut3*^*CreER*^ mouse. All the cKO mice were crossed in mixed genetic background and their littermates were selected as control for each experiment. No obvious difference in body size and weight were noticed in the littermates. Tamoxifen (Sigma, 20mg/mL in corn oil) was injected intraperitoneally (i.p.) into the mice with *CreER* background at P8 with a dose of 3mg/40g (tamoxifen/body weight). The mice were injected once a day for 3 days according to the references. The experimental procedures on mice were approved by the Institutional Animal Care and Use Committee of Tsinghua University.

### Nail auditory brainstem response (nABR)

Mice with either sex were anesthetized (i.p.) with 0.4% pentobarbital sodium in saline (0.2mL/10g, volume/body weight). Body temperature was maintained at 37°C by a heating pad during the whole experiment. After the skin on the vertex was removed, the skull was exposed and nailed with a stainless-steel screw (M1.4*2.5) but not puncturing the dura. The recording electrode was connected to the screw by a silver wire with a diameter of 0.1 mm. Other operations were similar to regular ABR procedure. The reference electrode and the ground electrode were inserted subcutaneously at the pinna and the groin respectively. The mice harboring a bone screw in Type-A implantation best-exhibited the ultrasonic responses (Fig. S1), which was used in this study. The ABR data were collected with ∼ 200-kHz sampling rate by an RZ6 workstation controlled by a BioSig software (Tucker-Davis Technologies, Alachua, FL). Clicks and 4-16 kHz pure-tone bursts were generated by a TDT MF1 closed-field magnetic speaker while a TDT EC1 (Coupler Model) electrostatic speaker was used for generating high frequencies (32-80 kHz). For the sound stimulation, 0.1-ms duration of click stimulus and 5ms duration with 0.5 ms rise-fall time of tone bursts were delivered at 21 Hz, with intensities from 90 to 10 dB SPL in 10 dB steps. Upon each acoustic stimulation with defined frequency and intensity level, the responses were sampled 512 times repeatedly and then averaged. The lowest stimulus sound level at which a repeatable wave 1 could be identified was defined as the threshold as reported (45).

### Acoustic-cue associated freezing behavior

The male mice were used. The mouse locomotion in an operant box (cubic, 30×30×30 cm) or an activity box (cylindrical, diameter of 35 cm and height of 30 cm) was monitored by an infrared camera with an infrared light as the light source, which was performed in a sound proof chamber (Shino Acoustic Equipment Co., Ltd, Shanghai, China). Each mouse was allowed to freely explore the operant box for 30 min before the sound-associated footshock training. During the training, an acoustic cue of 10 s containing 50 ms pure tone (16 kHz or 63 kHz) at 50 ms interval was played, and electrical shocks of 1 s at current magnitude of 0.6 mA was given to the mouse at the 5^th^ s and 10^th^ s. In the operant box, the electrical shocks were delivered by the metal grid floor powered by an electrical stimulator (YC-2, Chengdu Instrument Inc., Chengdu, China), and the acoustic cues were given by a free-field electrostatic speaker ES1 placed 15 cm above the floor and powered by an RZ6 workstation and a BioSig software (Tucker-Davis Technologies, Alachua, FL). The cue was given every 3 min and repeated for 10 times before the trained mouse was put back to the home cage. After 24 hours, the trained mouse was transferred in an activity box to test the freezing behavior. In the activity box, the same ES1 speaker was placed 15 cm above the chamber floor to generate the 16 kHz or 63 kHz acoustic cues of 10 s duration (identical to the training cues) and the cues were given at least 5 times during the test procedure. As calibrated, the sound intensity on the arena floor was from 70 dB SPL to 90 dB SPL that is in the range of mouse hearing threshold (Fig. S3).

### Open field test

The male mice at 1 month age were used. The mice were put in a cylindrical box with diameter of 35 cm and height of 30 cm that was a new place to them. The locomotion was recorded with an infrared camera and illuminated with an infrared LED. The distance a mouse travelled in the first 5 min was calculated with MATLAB 2014b (MathWorks, Natick, MA).

### Immunostaining

The mice were selected for immunostaining at indicated ages. After anesthesia with Avertin (30 mg/mL in saline, 0.12-0.15 mL/10g for mice), the mouse was perfused with ice-cold Phosphate Buffered Saline (PBS) and sacrificed by decapitation and the inner ears were dissected from the temporal bone. Then the inner ears were fixed by fresh 4% Paraformaldehyde (PFA) (DF0135, Leagene, Anhui, China) in PBS for 12-24 h at 4 °C. After fixation, the inner ears were washed with PBS for three times (10 min for each time), and then were treated in 120 mM EDTA decalcifying solution (pH 7.5) for 24 hours at room temperature (RT, 20 - 25 °C) followed by PBS washing. The cochlear coils were finely dissected from the inner ears in PBS and blocked in 1% PBST (PBS + 1% Triton X-100 (T8787, Sigma-Aldrich, St. Louis, MO)) solution with 5% BSA (A3059, Sigma-Aldrich, St. Louis, MO) at RT for 1 h. The cochlear tissues were then incubated in 0.1% PBST 5% BSA solution with MYO7A antibody (1:1000, Cat.25-6790, Proteus Biosciences Inc., Ramona, CA) overnight at 4 °C and washed with 0.1% PBST for three times at RT. The tissues were incubated with secondary antibody (Invitrogen anti-Rabbit Alexa Fluor 647, 1:1000, A21244; Invitrogen Alexa Fluor 488 Phalloidin, 1:1000, Cat. A12379) and 1:1000 DAPI in 0.1% PBST 5% BSA solution at RT for 2-4 hours. Tissues were washed with 0.1% PBST three times and mounted by ProLong Gold Antifade Mountant (Cat. P36930, Life Technology, Rockville, MD). The photos of fluorescent immunostaining pattern were collected by an A1/SIM/STORM confocal microscope (A1 N-SIM STORM, Nikon, Japan). The whole-view photos of the cochlear tissues were stitched by Photoshop software (version 9.3.1, Bitplane, Oxford instruments, Abingdon, England). The immunostaining procedure of cochlear tissues from *Piezo2-GFP* mice was slightly changed based on the protocol above. For fixation, the inner ears were perfused by 2% fresh PFA and incubated at RT for 30-45 min without shaking. For blocking, the cochlear tissues were treated in 0.5% PBST solution with 4% BSA at RT for 2 h with slow shaking. The tissues were incubated overnight at 4 °C with the primary antibody (Rabbit anti-GFP; 1:500, Cat. A-11122, ThermoFisher Scientific, Waltham, MA) that was made in 0.5% PBST with 1% BSA, and then washed with 0.1% PBST 3 times. Then the cochlear tissues were incubated in the secondary antibody (Invitrogen anti-Rabbit Alexa Fluor 647, 1:1000, Cat.A21244; Invitrogen Alexa Fluor 568 Phalloidin, 1:1000, Cat. A12380) was made in 0.5% PBST solution. Each incubation was shaken slowly.

### Inner ear clarification

We use PEGASOS method for inner ear clarification as previously reported (32). Mice were anesthetized by 0.4% pentobarbital sodium with an i.p. injection, followed by transcardiac perfusion with ice-cold 0.01M PBS to wash out blood and then with 4% PFA. Inner ear was dissected in 4% PFA and fixed for 12h at room temperature. After that, the following steps were performed in 37°C shaker. Inner ear was immersed in 0.5M EDTA for 2 days with daily change for decalcification and ddH_2_O for 2h to wash out remaining salt. Next, inner ear was immersed in 25 % Quadrol (diluted with H_2_O to a final concentration of 25% v/v, 122262, Sigma-Aldrich) for 2 days with daily change and 5% ammonium solution (diluted with H_2_O to a final concentration of 5% v/v, 105432, Sigma-Aldrich) for 6h to decolorize. Then, ammonium solution was washed out with PBS for 30min, followed by immunostaining steps. Inner ear was firstly immersed in blocking solution (4% BSA (V900933, VETEC) in 0.5% PBS-Triton X-100 (T8787, Sigma-Aldrich)) for 1 day, followed by Myo7a antibody (1:800, Rabbit, 25-6790, Proteus Biosciences) in blocking solution for 2 days with daily change. Then, 1^st^ antibody was washed out with 0.5% PBS-Triton X-100 for 1 day. Goat anti-Rabbit secondary antibody Alexa Fluor 488 (1:800, A11008, Thermo-Fisher) in blocking solution was used to incubate for 2 days with daily change, also followed by washing out with PBS for 1 day. After that, delipidation was performed with inner ear immersed in 30% tert-butanol (diluted with H_2_O, 360538, Sigma-Aldrich) for 4h, 50% tert-butanol for 6h, 70% tert-butanol for 1 day. Then, inner ear was immersed in tB-PEG (70% v/v tert-Butanol, 27% v/v PEG methacrylate Mn 500 (PEGMMA500) (409529, Sigma-Aldrich) and 3% w/v Quadrol) for 2 days with daily change for delipidation and BB-PEG (75% v/v benzyl benzoate (BB) (W213802, Sigma-Aldrich), 22% v/v PEGMMA500 and 3% w/v Quadrol)) for 2 days with daily change for clearing. Clarified inner ear was imaged with light-sheet microscope (Zeiss, Lightsheet Z.1) using 5× objective lens.

### RNAscope detection

After anesthesia with Avertin, transcardiac perfusion was done in P21 mice with ice-cold DEPC-PBS and then with 4% PFA (dilute from 16% PFA, DF0131, Leagene, Anhui, China) in DEPC-PBS. Then the temporal bones were dissected and fixed in fresh 4% PFA in DEPC-PBS at 4°C for 12h. After the post-fixation, the cochleae were decalcified by incubating at 120mM EDTA decalcification solution at RT for 48-60 h. Then the cochleae were dehydrated in 15% sucrose solution in DEPC-PBS for about 30 min at 4°C and in 30% sucrose solution in DEPC-PBS for about 2h until the cochleae sunk to the bottom of the tubes. After that the cochleae were embedded in O.C.T (4583, Tissue-Tek, Torrance, CA) and stored at -80°C. The embedded tissues were sliced into 14-μm sections (CryoStarTM NX50, ThermoFisher Scientific, Waltham, MA) and stored at -20°C no more than 8h before RNAscope detection. The *Piezo2* transcript detection was performed according to manufacturer’s instructions using RNAscope detection kit (323100, ACDBio, Newark, CA). We had taken a hard time to optimize the procedure because the organ of Corti was difficult to keep the original shape and stick on the glass slide after stringent treatments. This procedure was also tested by control probes (Fig. S4E). The probe of *Piezo2* (439971, ACDBio, Newark, CA) was validated by staining in DRG tissue slice (Fig. S4F). The sections of organ of Corti from control and *Piezo2*-cKO mice were placed on a same glass slide to encounter the same RNAscope procedure and imaging conditions.

### Hemicochlear imaging

Mice at 1-month age were anesthetized by isoflurane and sacrificed, and then their cochleae were dissected out in the dissection solution containing (in mM): 5.36 KCl, 141.7 NaCl, 1 MgCl_2_, 0.5 MgSO_4_, 0.1 CaCl_2_, 10 H-HEPES, 3.4 L-Glutamine, 10 D-Glucose (pH 7.4, Osmolarity at 290 mmol/kg). Immersed in the cutting solution containing (in mM): 145 NMDG-Cl, 0.1 CaCl_2_, 10 H-HEPES, 3.4 L-Glutamine, 10 D-Glucose (pH 7.4, Osmolarity at 290 mmol/kg), the cochlea was glued on a metal block with Loctite 401 and cut to 2 halves by a vibratome (VT1200S, Leica, Wetzlar, Germany) with FREQ index at 7, Speed index at 50. The section plane should be parallel to the modiolus to minimize the damage on tissue. The hemicochlea was transferred into a recording dish, and glued on the bottom, and loaded with 25 μg/mL Fluo-8 AM (Invitrogen, Waltham, MA) in the recording solution. After 10-min incubation at RT in a dark box, the dye-loading solution was replaced by the dye-free recording solution containing (in mM): 144 NaCl, 0.7 Na_2_PO4, 5.8 KCl, 1.3 CaCl_2_, 0.9 MgCl_2_, 10 H-HEPES, 5.6 D-Glucose (pH 7.4, Osmolarity at 310 mmol/kg). An upright microscope (BX51WI, Olympus, Tokyo, Japan) equipped with 60× water immersion objective (LUMPlanFL, Olympus, Tokyo, Japan) and an sCMOS camera (ORCA Flash 4.0, Hamamatsu, HAMAMATSU-SHI, Japan) was used for calcium imaging, controlled by MicroManager 1.6 software (46) with a configuration of 4×4 binning, 100-ms exposure time, and 2-s sampling interval. To keep the best performance of the hemicochlea preparations, the whole procedure from cutting to imaging was finished within 15 min to guarantee the best appearance of tissue samples. As control experiments, 0.1 mM Ca^2+^ (to keep tip link structure) perfusion abolished the ultrasonic stimulation evoked Ca^2+^ signal, and 100 μM ATP perfusion induced strong Ca^2+^ response (∼20%), in the OHCs of the hemicochleae.

### Ultrasound generation and delivery *ex vivo*

A customized 80-kHz ultrasound transducer with diameter of 27 mm was powered by a radio-frequency amplifier (Aigtek, ATA-4052, China) integrated with a high-frequency function generator (Rigol, DG1022U, China). The 80-kHz transducer was chosen because its size is small enough to be assembled (the lower the frequency, the larger the size) and 80 kHz is a physiological frequency to mice. For calibration, a high-sensitivity hydrophone (Precision Acoustics, United Kingdom) was positioned directly above the vibration surface. Transducer outputs were calibrated in a tank filled with deionized, degassed water under free-field conditions. To stimulate hemicochlea, the transducer was tightly fixed at the bottom of recording dish with ultrasound gel in between. The distance between the tissue and ultrasound transducer is less than 5 mm. For the 80-kHz ultrasonic stimulation, a single pulse of 100 ms was applied, with calibrated intensities at 8.91 W/cm^2^ I_SPTA_. The ultrasound energy received by the tissue preparation was stable and homogeneous, as shown by calibrated intensities covering the whole bottom of the recording dish (Fig. S7).

### Low-frequency fluid-jet stimulation to hemicochlea

Fluid-jet configuration was used as previously reported (47). Briefly, a 35-mm diameter circular piezoelectric ceramic was sealed in a self-designed mineral oil tanker. An electrode with 5-10 μm diameter tip filled with recording solution (144 NaCl, 0.7 Na_2_PO_4_, 5.8 KCl, 1.3 CaCl_2_, 0.9 MgCl_2_, 10 H-HEPES, 5.6 D-Glucose in mM, pH 7.4, Osmolarity at 310 mmol/kg) was mounted into the tanker and transmitted the pressure wave to the hair bundle of an OHC in hemicochlea samples. The circular piezoelectric ceramic was driven by a sinusoidal voltage fluctuation generated from a patch-clamp amplifier (EPC10 USB, HEKA Elektronik, Lambrecht/Pfalz, Germany) and amplified at 20 folds with a custom high-voltage amplifier. The 100-ms sinusoidal stimulation was given at frequency of 2000 Hz and amplitude of 6.5 V.

### Single-cell Ca^2+^ imaging and whole-cell electrophysiology

HEK293T cells were plated onto 8-mm round glass coverslips, which were coated with poly-D-lysine and placed in 48-well plates. 400 ng of plasmids were transiently transfected into HEK293T cells using lipofectine 2000 (Life Technologies). GCaMP6 was expressed to monitor the Ca^2+^ response. After 24h transfection, the HEK293T cells were imaged for Ca^2+^ signals by an upright microscope (BX51WI, Olympus, Tokyo, Japan) equipped with 60× water immersion objective (LUMPlanFL, Olympus, Tokyo, Japan) and an sCMOS camera (ORCA Flash 4.0, Hamamatsu, HAMAMATSU-SHI, Japan), controlled by MicroManager 1.6 software (46) with 50-ms exposure time and 1-s sampling interval. HEK293T cells were recorded using whole-cell patch-clamp as previously described (48). All experiments were performed at room temperature (20-25°C). Briefly, the coverslip with cultured cells was transferred into a recording chamber with recording solution containing (in mM): 144 NaCl, 0.7 NaH_2_PO_4_, 5.8 KCl, 1.3 CaCl_2_, 0.9 MgCl_2_, 5.6 glucose, and 10 H-HEPES (pH 7.4). The cells were imaged under an upright microscope (BX51WI, Olympus, Tokyo, Japan) with a 60× water-immersion objective and an sCMOS camera (ORCA Flash4.0, Hamamatsu, Hamamatsu City, Japan) controlled by MicroManager 1.6 software (46). Patch pipettes were made from borosilicate glass capillaries (BF150-117-10, Sutter Instrument Co., Novato, CA) with a pipette puller (P-2000, Sutter) and polished on a microforge (MF-830, Narishige, Tokyo, Japan) to resistances of 4-6 MΩ. Intracellular solution contained (in mM): 140 KCl, 1 MgCl_2_, 0.1 EGTA, 2 Mg-ATP, 0.3 Na-GTP, and 10 H-HEPES, pH 7.2). The cells were recorded with a patch-clamp amplifier with a holding potential of –70 mV (EPC 10 USB and Patchmaster software, HEKA Elektronik, Lambrecht/Pfalz, Germany). The liquid junction potential is not corrected in the data shown. As measured, the pipette with CsCl intracellular solution had a value of +4 mV in regular recording solution.

Mechanical stimulation utilized a fire-polished glass pipette (tip diameter 3–4 mm) positioned at an angel of 80 relative to the cell being recorded as described (48). The probe was displaced by a piezoelectric actuator (P-601.1SL, Physik Instrumente, Karlsruhe, Germany) and driven by a piezoelectric crystal microstage (E625 LVPZT Controller/Amplifier, Physik Instrumente, Karlsruhe, Germany). The probe velocity was 1 μm/ms during the upward and downward movement, and the stimulus was kept constant for 100 ms. A series of mechanical steps in 1 μm increments was applied every 5–10 s.

### Data analysis

Each experiment contained at least 3 biological replicates. Data were managed and analyzed with MATLAB 2014b (MathWorks, Natick, MA), MicroManager 1.6 software (46), Excel 2016 (Microsoft, Seattle, WA), Prism 6 (GraphPad Software, San Diego, CA), and Igor pro 6 (WaveMetrics, Lake Oswego, OR). All data are shown as mean ±SD, as indicated in the figure legends. We used two-tailed *t*-test for one-to-one comparison or one-way ANOVA for one-to-many comparison to determine statistical significance (*p<0.05, **p<0.01, ***p<0.001, ****p<0.0001). N numbers are indicated in the figures. For Animal tracing and locomotion evaluation, videos of mouse locomotion in open-field, foot-shock and pup-retrieval test were analyzed by MATLAB software and EthoVision XT software (v11.5, Noldus, Wageningen, Netherland). The center of mice was used to draw locomotion trace. To show the speed information, the locomotion trace was dotted every 0.5 s. For footshock behavior analysis, freezing time percentage of pre-cue (30s before conditional stimulus) and post-cue (30s after conditional stimulus) were analyzed to compare the effect of sound induced freezing. For Ca^2+^ data analysis, to extract fluorescence signals, we visually identified the regions of interest (ROIs) based on fluorescence intensity. To estimate fluorescence changes, the pixels in each specified ROI were averaged (F). Relative fluorescence changes, ΔF/F_0_ = (F-F_0_) / F_0_, were calculated as Ca^2+^ signals. The hemicochlear imaging data were analyzed offline by Micromanager software and Excel software. The ROI was drawn to cover each hair cell. The fluorescence intensity of ROI was normalized to its value in the frame right before the stimulation.

## Acknowledgments

We thank Drs Xiaoqin Wang, Xiaoke Chen, Lei Song, Xin Liang, and members of Xiong laboratory for helpful discussions and critical proof-reading of this manuscript, thank the Imaging Core Facility, Technology Center for Protein Sciences at Tsinghua University for assistance of using imaging instruments and software, thank Si Li of Dr. Qingfeng Wu laboratory for help with RNAscope experiment, thank Lili Niu and Xudong Shi of Dr. Hairong Zheng laboratory at Shenzhen Institutes of Advanced Technology, Chinese Academy of Sciences for manufacturing the high-frequency amplifier and ultrasonic transducers, and thank Dr. Qiuying Chen of Dr. Guoxuan Lian laboratory at Institute of Acoustics, Chinese Academy of Sciences for manufacturing the ultrasonic transducers, and thank Dr. Guangzhen Xing at National Institute of Metrology for calibrating the ultrasonic transducers and the high-frequency amplifier. This work was supported by the National Natural Science Foundation of China (31522025, 31571080, 81873703, 3181101148, and 31825014), Beijing Municipal Science and Technology Commission (Z181100001518001), National Key Scientific Instrument and Equipment Development Project (81527901) and a startup fund from the Tsinghua-Peking Center for Life Sciences, W.X. is a CIBR cooperative investigator (2020-NKX-XM-04) funded by the Open Collaborative Research Program of Chinese Institute for Brain Research.

